# An unusual ring pattern in the Rosβ lanthipeptide of the two-component lantibiotic roseocin

**DOI:** 10.1101/2025.10.27.684949

**Authors:** Emily K. Desormeaux, Lingyang Zhu, Youran Luo, Dipti Sareen, Wilfred A. van der Donk

## Abstract

Two-component lantibiotics comprise two post-translationally modified peptides that synergistically exert antimicrobial activity. Most known two-component lantibiotics are made up of a structurally conserved α-lanthipeptide that binds to the peptidoglycan precursor lipid II and a β-lanthipeptide that is believed to interact with the α-peptide-lipid II complex to form pores in the membranes of susceptible bacteria. A few two-component lantibiotics including roseocin do not follow this general scheme and act by different, currently unresolved mechanisms. An important first step in studying this group of lanthipeptides is determination of their chemical structure. Roseocin’s β-peptide (Rosβ) is formed by the RosM lanthipeptide synthetase from the RosA1 precursor peptide. RosM carries out nine dehydrations of Ser and Thr residues to the corresponding dehydroamino acids followed by six Michael-type additions of the thiols of Cys residues to a subset of the dehydroamino acids forming six thioether crosslinks. Sequence alignment with structurally characterized lanthipeptides does not allow prediction of its ring pattern. In this study, Rosβ was produced in *Escherichia coli* and its ring pattern was established by multi-dimensional NMR spectroscopy. The stereochemistry of the lanthionine and methyllanthionine residues was determined by Marfey’s analysis with authentic standards. Rosβ is shown to have a unique ring pattern amongst previously characterized lanthipeptides.

**TOC graphic:** 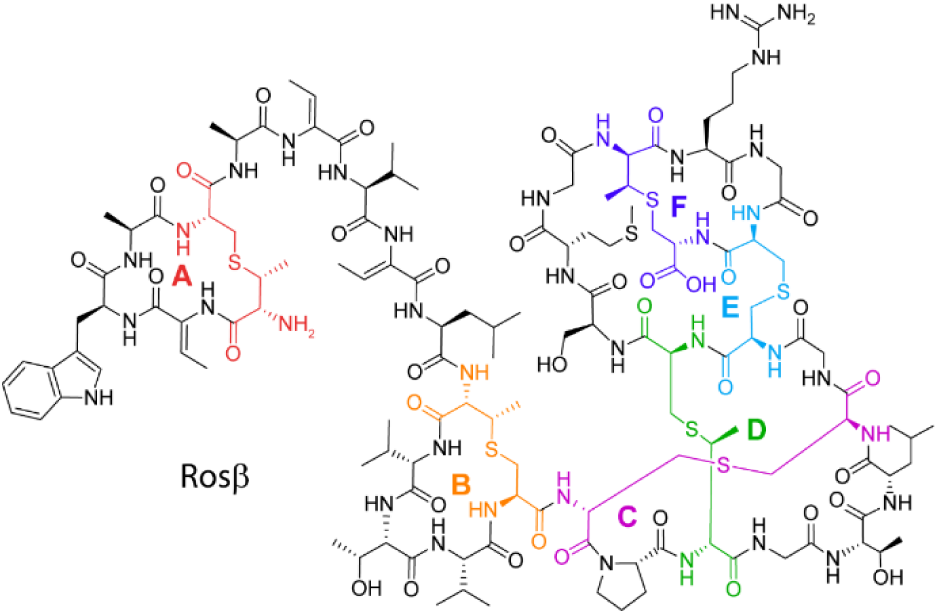

---

Lanthipeptides are a group of peptidic natural products containing thioether crosslinks that are introduced by post-translational modification.^1^ Two-component lantibiotics form a subgroup of these natural products that comprise two different post-translationally modified lanthipeptides that display synergistic antimicrobial activity.^2,3^ Thus far, all reported two-component lantibiotics have been members of class II lanthipeptides, for which a single enzyme (generically termed LanM) introduces multiple thioether macrocycles by first dehydrating selective Ser and Thr residues in their precursor peptides by phosphorylation of the alcohol side chains followed by phosphate elimination.^4^ This process results in the formation of the dehydroamino acids dehydroalanine (Dha) from Ser and dehydrobutyrine (Dhb) from Thr (Figure 1A). Subsequently, in a different active site, the same enzyme catalyzes the Michael-type addition of Cys residues to the Dha and Dhb residues to form lanthionine (Lan) and methyllanthionine (MeLan) crosslinks, respectively.^5^ These modifications take place in the C-terminal part of the precursor peptide called the core peptide; the N-terminal part of the precursor peptide, termed the leader peptide, serves as a recognition element for the post-translational modification enzymes and is removed by a protease in what is usually the last step of maturation.^6^ The biosynthesis of several two-component lantibiotics has been studied demonstrating that the two individual subunits are produced either by a singular LanM enzyme, capable of modifying both peptides,^7–9^ or by two distinct LanM enzymes, each modifying one of the two precursor peptides.^10–12^

**Figure 1.**
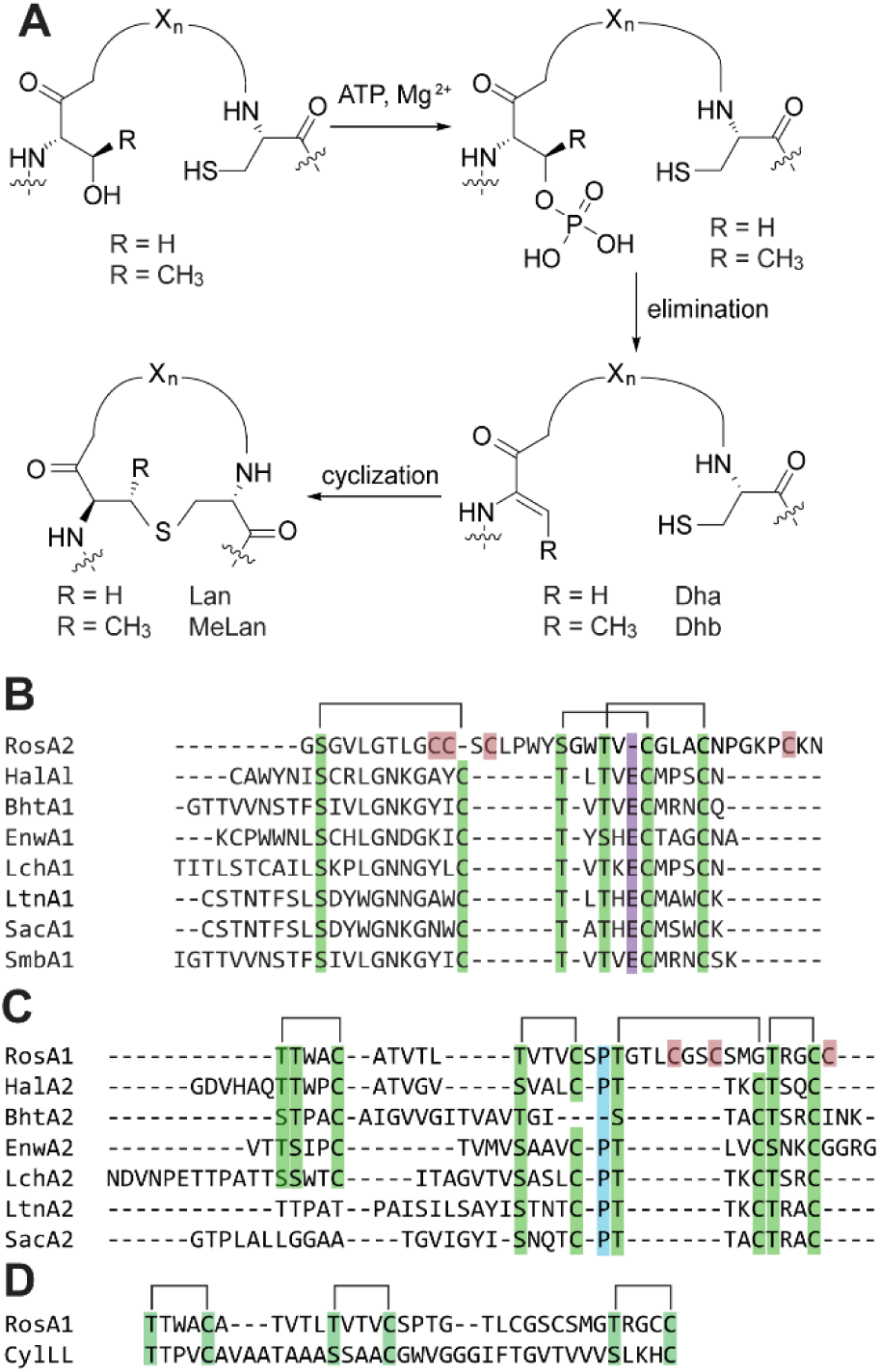
Lanthipeptide biosynthesis and structures of selected two-component lantibiotics. **A)** Post-translational modification process for class II lanthipeptides that results in Lan and MeLan. For clarity, only products with DL sterochemistry are shown. **B)** Sequence alignment of the predicted core regions of the predicted α-peptides of a group of structurally similar two-component lantibiotics. The glutamate residue, absent in RosA2 and critical for lipid-II binding and antimicrobial activity in mersacidin-like peptides, is shown in purple. The additional Cys residues in RosA2 that are not conserved in the other peptides are highlighted in red. These additional Cys residues are conserved in roseocin analogs.^23^ The ring pattern shown above the sequences is for a subset of known α-peptides that have conserved Cys and Ser/Thr residues (green) that form the indicated crosslinks. These ring-forming residues and the crosslinks they generate are not present in all family members. For RosA2, three Cys residues **C)** Sequence alignment of the predicted core regions of the partner β-peptides. The additional Cys residues in RosA1 that are not conserved in the other peptides are highlighted in red. Rings formed in other β-peptides from conserved Cys and Ser/Thr residues (green) are indicated above the alignments; roseocin β does not contain all ring-forming residues of other members, and has additional ring-forming residues that do not align with other members. In blue is a Pro that is often conserved in β-peptides. **D)** Alternative manual alignment of the RosA1 core peptide with the CylL_L_ core peptide. Hal, haloduracin from *Bacillus halodurans*;^11,24^ Bht, peptides from *Streptococcus rattus* strain BHT;^25^ Enw, enterocin W from *Enterococcus faecalis* NKR-4-1;^26^ Lch, lichenicidin from *Bacillus licheniformis* DSM 13 and VK21;^27–29^ Ltn, lacticin 3147 from *Lactococcus lactis* DPC3147;^30^ Sac, staphylococcin C55 from *Staphylococcus aureus* C55;^31^ Smb, peptides from *Bacillus licheniformis* DSM 13.^32^

The mechanism of action of most of the studied two-component lantibiotics involves a 1:1 ratio of α- and β-peptides.^13–15^ The α-peptides of the majority of these natural products are structurally highly conserved (Figure 1B) and bind to the peptidoglycan biosynthetic precursor lipid II using a sequence motif containing a highly conserved glutamate residue, analogous to that found in the single-component lanthipeptide mersacidin (Figure 1A).^16–19^ The partner β-peptides have less sequence similarity than their α-counterparts, but still often have conserved ring patterns in characterized peptides (Figure 1C). Whereas the constituent peptides of most reported two-component lantibiotics contain similar ring patterns as those shown in Figure 1, a handful of two-component lantibiotics have been reported that do not have this general architecture. These are cytolysin produced by *Enterococcus faecalis* and the structurally related bibacillins,^7,20^ bicereucin produced by *Bacillus cereus* SJ1,^9^ and roseocin encoded by *Streptomyces roseosporus* NRRL11379/15998 and *Streptomyces* sp. ADI96-02 (Figure S1).^8,21,22^ Roseocin is the first two-component lantibiotic found in the genomes of members of Actinobacteria and the ring patterns of its two constituents (Rosα and Rosβ) have not yet been determined. Several analogs of roseocin have also been identified bioinformatically.^23^

The roseocin biosynthetic gene cluster (BGC) encodes the two precursor peptides (RosA1 and RosA2), one LanM enzyme (RosM), a transporter (RosT_p_), likely immunity proteins (RosEFG), and a protein of unknown function (Figure 2).^8^ A previous investigation showed that co-expression of RosM with each of the individual precursor peptides separately in *E. coli* resulted in modified peptides that when purified displayed strong synergistic antibacterial activity after leader peptide removal.^8^ Furthermore, a study on the native producer *S. roseosporus* NRLL15998 detected nine-fold dehydrated Rosβ, consistent with the product made in *E. coli;* Rosα was not detected in that study.^21^ Rosα is generated from the RosA2 precursor peptide, and Rosβ is produced from the RosA1 precursor peptide. Whereas the core peptide of RosA2 has some sequence homology to the precursor peptides of other α-peptides (but lacks the lipid II binding motif, Figure 1B), the core peptide of the RosA1 precursor is divergent from other β-peptides. The constellation of Cys, Ser, and Thr residues at the N- and C-termini may result in three rings that are at similar positions as in other β-peptides. That would still leave three more Cys residues that are not present in those peptides (Figure 1C). In this study, we determined the ring pattern and stereochemistry of the Rosβ peptide using a combination of NMR spectroscopy, Marfey’s analysis, and mass spectrometry (MS). In parallel, we also attempted structural elucidation of the Rosα peptide, but poor solubility in a variety of solvents and poor signal dispersity suggesting aggregation, precluded NMR characterization (see Experimental Section).

**Figure 2.**
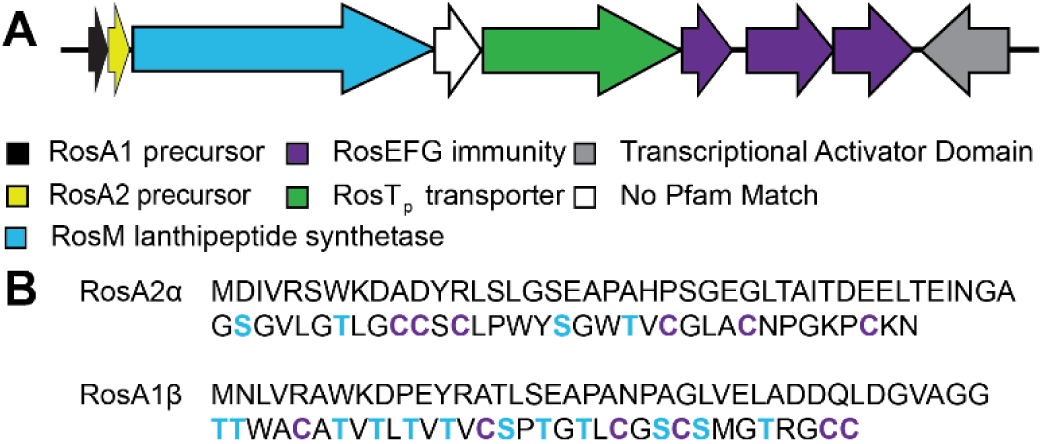
Biosynthetic gene cluster of roseocin. **A)** BGC diagram with genes involved in the biosynthesis of roseocin. The BGC includes genes encoding the RosA1 and RosA2 precursor peptides, a gene encoding the RosM lanthipeptide synthetase, a gene encoding a transporter/protease, three genes likely involved in self-immunity, and two additional genes that are not involved in the chemistry of biosynthesis. **B)** Precursor sequences for Rosα and Rosβ. The predicted cleavage sites between leader and core peptides are indicated by a line break. Potentially dehydrated residues (Ser/Thr) in the putative core peptide are indicated in blue and Cys residues in purple.

## RESULTS AND DISCUSSION

### Sequence alignment of roseocin precursor peptides

Multiple sequence alignment of both the α and β core peptides of selected two-component lantibiotics was performed (Figure 1). The Rosα peptide aligned well with other α-peptides with the exception of two regions, one located within the first large ring of other α-peptides and another located at the C-terminus of the peptide. These regions of Rosα also contain two additional Cys residues that were identified to be involved in disulfide formation.^8^ Thus, Rosα contains four thioether rings as opposed to three in known α-peptides. Most notably, this alignment shows that Rosα is lacking the highly conserved glutamate residue that is found within the lipid II binding domain of mersacidin (Figure S1) that is essential for its bioactivity.^18,33,34^ In fact, the Rosα core peptide sequence does not contain any Glu or Asp residues, suggesting a potential different molecular target or mechanism of action than that of other α-peptides.

The β-counterparts of two-component lantibiotics associated with the known α-peptides aligned less well with Rosβ, with three of the four conserved rings in other β-peptides potentially also present in Rosβ (Figure 1C). An additional manual alignment was also performed with CylL_L_ indicating a potentially alternative partially conserved ring pattern (Figure 1D). Cytolysin was the first studied two-component lanthipeptide and is composed of a large subunit, CylL_L_”, and a small subunit, CylL_S_” (Figure S1).^7,35,36^ Cytolysin has lytic activity against both bacterial and eukaryotic cell lines, a rare observation for lanthipeptides. Its activity against eukaryotic cells is directly responsible for poor disease prognosis for patients infected with cytolysin-producing *E. faecalis.*^37^ The various possible sequence alignments in Figure 1 highlight the need for a fully characterized structure of Rosβ.

### Production of Rosβ

We co-expressed a His-tagged version of the RosA1 precursor with untagged RosM in *Escherichia coli* as reported previously.^8^ After Ni^2+^-affinity purification, analysis by matrix-assisted laser desorption/ionization time-of-flight mass spectrometry (MALDI-TOF MS) indicated nine dehydrations (Figure S2). Several peptidases were utilized to attempt leader peptide removal including LahT150,^38^ the C39 peptidase domain of the peptidase-containing ATP-binding cassette transporter (PCAT) LahT, and endoproteinase AspN. Leader peptide removal with LahT150 was unsuccessful, potentially because the cleavage site is predicted to be right next to a methyllanthionine ring. To obtain a single Rosβ product, AspN was therefore utilized to remove a portion of the leader peptide (Figure S2). We then removed the remaining residues of the leader peptide using proteinase K.^8^ Using a similar procedure, we also produced Rosα, but its solubility properties were unsuitable for NMR analysis (see Experimental Section). Therefore, we concentrated on Rosβ for structural elucidation.

### Stereochemical analysis of Rosβ

After Ni^2+^-affinity purification, stereochemical analysis was performed by a modified Marfey’s analysis using liquid-chromatography coupled to mass spectrometry detection (LC-MS). This analysis using authentic standards as described previously^39^ indicated that the roseocin β-peptide contains DL-Lan, and both LL- and DL-MeLan (Figure 3). While Marfey’s analysis of these crosslinked bis-amino acids by LC-MS is not quantitative, the ratios of the peaks within each chromatogram are indicative of the ratios of stereoisomers within the peptide.^39^ The MeLan analysis indicates LL- and DL-products in an approximately one-to-three ratio, suggesting one LL-ring and three DL-rings. Using this information along with the precursor sequence, it was inferred that Rosβ contains four MeLan rings, and two Lan rings in the RosM-modified peptide.

**Figure 3.**
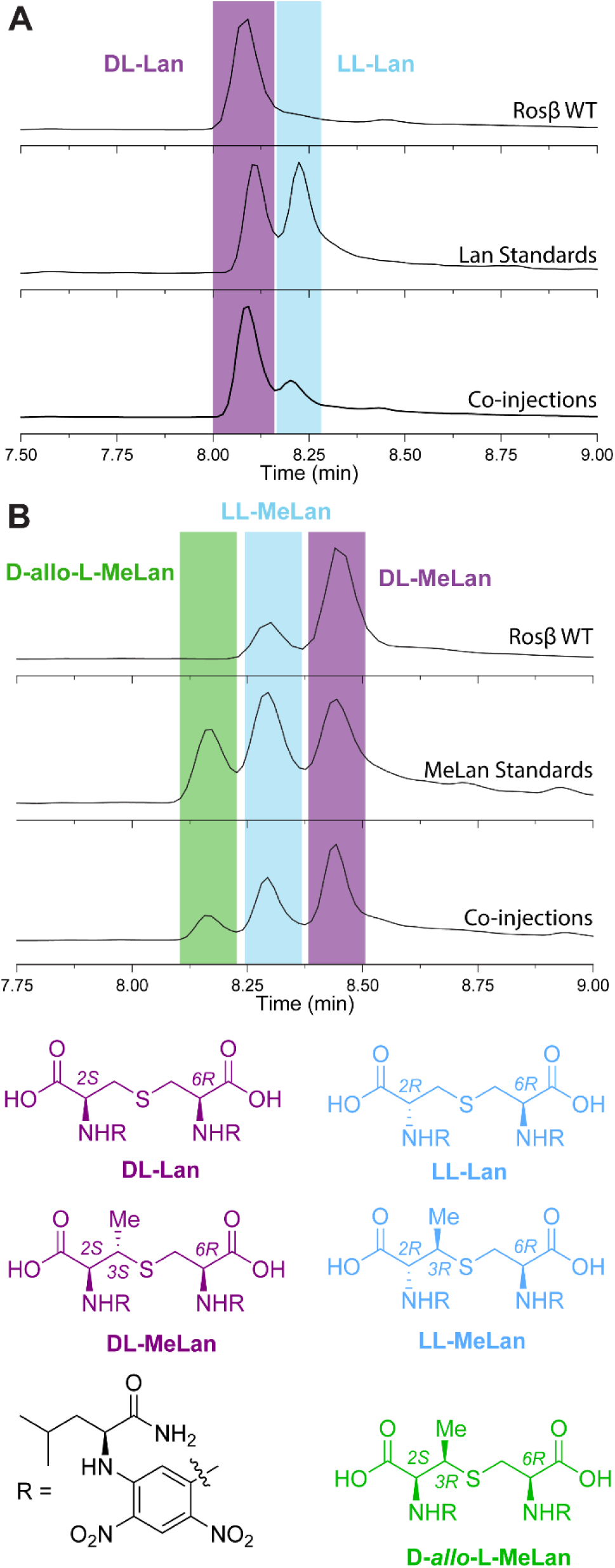
LCMS analysis of Marfey’s assay with RosM-modified RosA1β. **A)** Extracted ion chromatograms (EICs) of Marfey’s analysis of RosM-modified RosA1β (top), Lan standards (middle), and co-injections (bottom) analyzed by LC-MS for [M − H]^−^ m/z 795.2373. **B)** EICs of Marfey’s analysis of RosM-modified RosA1β (top), MeLan standards (middle), and co-injections (bottom) analyzed by LC-MS for [M − H]^−^ m/z 809.2530. Coloring of each peak: purple, DL stereochemistry; blue, LL stereochemistry; green, D-*allo-*L-MeLan stereochemistry. The products of the Marfey-derivatization are shown.

### NMR analysis of Rosβ

NMR data on Rosβ was obtained in a mixture of acetonitrile and water (1:1) using peptides with natural isotopic abundance. All peptide spin systems were assigned using a TOCSY spectrum (Figure 4). Positions of the amino acid assignments within the Rosβ sequence were determined by nuclear Overhauser effect spectroscopy (NOESY) using sequential amide proton assignments, as well as nuclear Overhauser effects (NOEs) between NH residues and the αH or side chain proton signals of their neighboring residues. These spectra were used collectively to assign all spin systems (Table 1). Amino acid assignments indicated that Thr1, Thr2, Thr7, Thr9, Thr11, Ser16, Thr18, Ser24, and Thr29 were all dehydrated by RosM, while residues Thr13, Thr20, Ser26 escaped dehydration in the RosM-modified Rosβ. These observations were consistent with previously published tandem MS data.^8^ The NMR data also indicated that thioether crosslinks were formed at Dhb1, Dhb11, Dha16, Dhb18, Dha24, and Dhb29. We will use the notations Ala16-Cys22 to indicate a lanthionine linkage between the former Ser16 and Cys22; for MeLan we will use the notation Abu1-Cys5 (*vide infra*) to indicate a linkage between aminobutyric acid (Abu) 1 (derived from Thr1) and Cys5.

**Figure 4.**
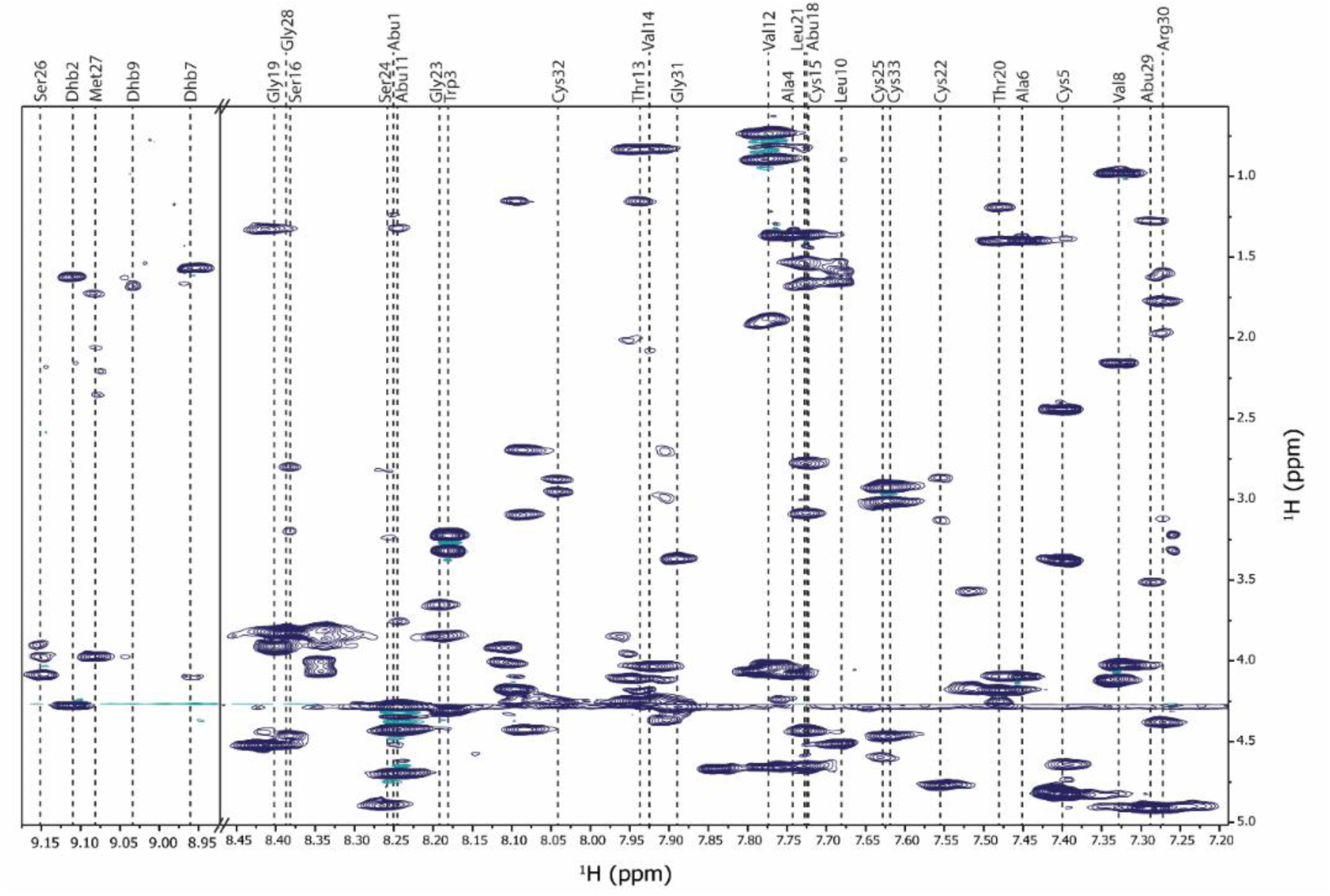
TOCSY spectrum of Rosβ in 50% ACN-d_3_/50%H_2_O showing the spin system of its component residues. The spectrum was taken with water suppression and a 70 ms relaxation time. Full spectrum without any axis breaks is provided in the SI (Figure S3). An overlay of TOCSY and NOESY spectra is shown in Figure S4.

**Table 1.** Chemical shift assignments for Rosβ in 50% ACN-d_3_/50%H_2_O. Cys and Ser/Thr residues involved in a ring formation are represented in matching colors.

| # | AA ID | N-H | αH | βH | γH | δH | other |
| --- | --- | --- | --- | --- | --- | --- | --- |
| 1 | Abu | 8.25 | 4.70 | 3.55 | 1.24 |  |  |
| 2 | Dhb | 9.11 | N/A | 5.87 | 1.63 |  |  |
| 3 | Trp | 8.18 | 4.31 | 3.23, 3.32 | 2H:7.26 (s), N(1)H:10.00 (br, s),<br>4H:7.53 (d),<br>5H:7.03 (t), 6H:7.13 (d), 7H:7.41 (t) |  |  |
| 4 | Ala | 7.74 | 4.09 | 1.37 |  |  |  |
| 5 | Cys | 7.40 | 4.83 | 2.44, 3.39 |  |  |  |
| 6 | Ala | 7.45 | 4.10 | 1.40 |  |  |  |
| 7 | Dhb | 8.96 | N/A | 6.66 | 1.57 |  |  |
| 8 | Val | 7.33 | 4.03 | 2.15 | 0.98 |  |  |
| 9 | Dhb | 9.03 | N/A | 6.95 | 1.68 |  |  |
| 10 | Leu | 7.68 | 4.51 | 1.59, 1.65 | 1.53 | 0.82, 0.90 |  |
| 11 | Abu | 8.25 | 4.43 | 3.75 | 1.32 |  |  |
| 12 | Val | 7.78 | 4.04 | 1.88 | 0.89, 0.74 |  |  |
| 13 | Thr | 7.94 | 4.25 | 4.19 | 1.16 |  |  |
| 14 | Val | 7.92 | 4.04 | 2.08 | 0.83 |  |  |
| 15 | **Cys** | 7.73 | 4.44 | 2.78, 3.09 |  |  |  |
| 16 | **Ala** | 8.38 | 4.46 | 2.80, 3.20 |  |  |  |
| 17 | **Pro** | N/A | 4.66 | 2.30, 1.92 | 2.01 | 3.82, 3.64 |  |
| 18 | **Abu** | 7.72 | 4.63 | 3.70 | 1.36 |  |  |
| 19 | **Gly** | 8.40 | 3.91 |  |  |  |  |
| 20 | **Thr** | 7.48 | 4.25 | 4.18 | 1.19 |  |  |
| 21 | **Leu** | 7.73 | 4.65 | 1.66, 1.54 | 1.54 | 0.86, 0.82 |  |
| 22 | **Cys** | 7.55 | 4.77 | 3.13, 2.87 |  |  |  |
| 23 | **Gly** | 8.19 | 3.65<br>3.85 |  |  |  |  |
| 24 | **Ala** | 8.26 | 4.90 | 3.24, 2.83 |  |  |  |
| 25 | **Cys** | 7.63 | 4.63 | 2.93, 3.01 |  |  |  |
| 26 | **Ser** | 9.15 | 4.08 | 3.90, 3.99 |  |  |  |
| 27 | **Met** | 9.08 | 3.98 | 1.72, 2.06 | 2.35, 2.20 | $\epsilon\text{CH}_3$: 1.87<br>(s) | |
| 28 | **Gly** | 8.39 | 3.83 |  |  |  |  |
| 29 | **Abu** | 7.29 | 4.91 | 3.51 | 1.28 |  |  |
| 30 | **Arg** | 7.28 | 4.38 | 1.78, 1.61 | 1.97, 1.61 | 3.12 | $\epsilon\text{NH}$:7.13 |
| 31 | **Gly** | 7.89 | 3.37 |  |  |  |  |
| 32 | **Cys** | 8.04 | 4.26 | 2.95, 2.88 |  |  |  |
| 33 | **Cys** | 7.62 | 4.47 | 2.93, 3.01 |  |  |  |

Two assignments were made for residues eight through eighteen because of two conformations containing either a cis or trans isomer at Pro17, which were present in a ratio of 2:3 (cis:trans). Because the spectra were obtained in 50% acetonitrile, thus likely representing non-natural conformations, no three-dimensional modeling was performed. All reported assignments below correspond to the isomer with the trans conformation at Pro17.

The NOESY spectrum was used to determine the location of (Me)Lan within Rosβ, revealing six rings, two Lan and four MeLan. The (Me)Lan structures identified within the peptide will be termed rings A-F and were shown to be formed between the following residue pairs: Abu1-Cys5 (ring A, Figure S5), Abu11-Cys15 (ring B, Figure S6), Ala16-Cys22 (ring C, Figure S7), Abu18-Cys25 (ring D, Figure S8), Ala24-Cys32 (ring E, Figure S9), and Abu29-Cys33 (ring F, Figure S10). These rings were assigned through NOE interactions between the α, β, and γ (in the case of Abu) protons in the ring forming residues (indicated in red within Figures S5-S10). In some cases, weak NOE peaks were also observed between these side chain protons and the amide proton of a residue involved in ring formation; an example of such a correlation is shown for the C ring (Figure S7C), where NOE peaks were observed between the former Cys22 amide and the β protons of the former Ser16. The Dhb residues at positions 7 and 9 both had the *Z* stereochemistry, as expected for a LanM-catalyzed *anti*-elimination of phosphate (Figure 1A).

The described ring pattern observed in Rosβ (Figure 5A) has not been observed in other lanthipeptides thus far reported, and while the stereochemistry has not been determined for each individual ring, we hypothesize that ring A is the LL-MeLan ring, as it aligns well with previously characterized rings containing LL-stereochemistry, such as CylL_L_”,^7,40^ haloduracin β,^40^ geobacillin II,^41^ and carnolysin.^42^ All of these LL rings are formed from a precursor with the sequence DhxDhxXxxXxxCys (Dhx = Dha or Dhb; Xxx = other amino acids), which was shown to favor formation of the LL stereochemistry.^7,40^

**Figure 5.**
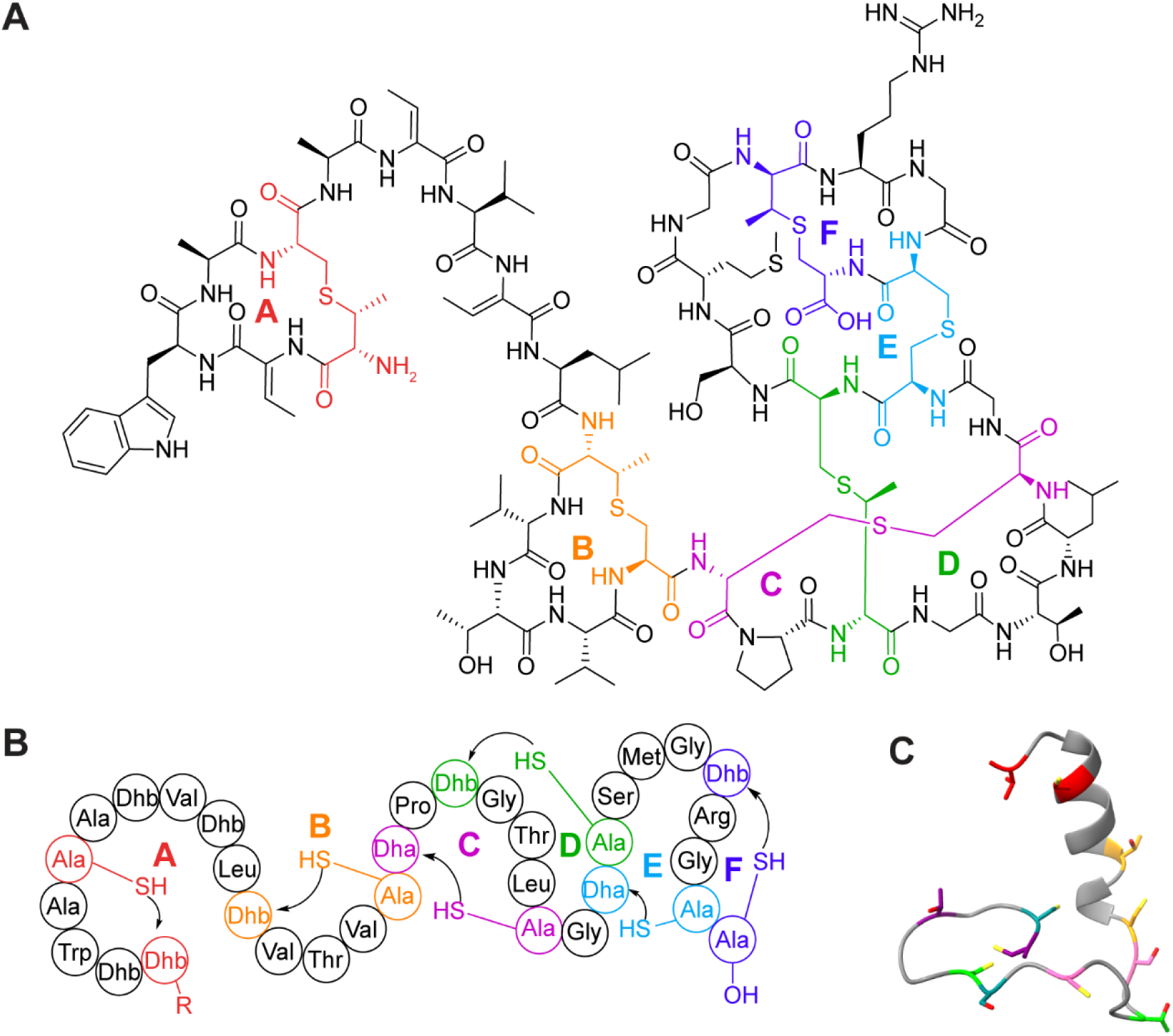
Structure of Rosβ. **A)** Chemical structure of Rosβ. The red MeLan most likely has the LL configuration based on sequence similarity to the ring at the N-terminus in other lanthipeptides for which LL-stereochemistry has been shown conclusively. **B)** Schematic representation of the formation of Rosβ. R = leader peptide. **C)** Lowest energy AlphaFold model of unmodified RosA1 core peptide suggesting an alpha helix in the N-terminal region (residues 1-15). Residues involved in ring formation are colored, matching those in Table 1 and panels A and B.

### Structure of roseocin β

Structural characterization of lanthipeptides is imperative for understanding the link between structure and bioactivity. Roseocin, which has antimicrobial activity against several clinically relevant bacterial strains,^8^ has novel precursor peptide sequences compared to other two-component lanthipeptides, suggesting a potential novel structure and mode of action. Indeed, roseocin’s β-peptide has a unique lanthipeptide structure containing six (Me)Lan rings, with the last four rings highly intertwined (Figure 5). The overall ring pattern resembles the large subunit of cytolysin (CylL_L_”) more than the ring patterns of other well-studied two-component lanthipeptides, with similar N-terminal A and B rings and the C-terminal F ring. However, the B ring in CylL_L_” has the LL-configuration whereas the B-ring in Rosβ has the DL-configuration. A recent bioinformatic analysis identified a number of BGCs encoding roseocin analogs.^23^ The NMR structure determined in this study shows that the A, B, D, E, and F rings are completely conserved in these analogs, whereas the residues forming the C-ring as well as the Pro in this ring are not always conserved.

The two N-terminal rings of Rosβ (A and B ring) show similarity in pattern and stereochemistry to both haloduracin β and lichenicidin β (Figure 6).^11,12,28,29^ For lichenicidin, the three dimensional structure was determined by NMR spectroscopy in acidic methanol (pH 3.5),^29^ showing an alpha helical structure from the start of the A-ring up until a Pro residue that is highly conserved in a number of lanthipeptides including Rosβ.^43^ The NMR structure of CylL_L_” in methanol also showed an α-helical structure for the N-terminus up to a stretch of three Gly residues. While the three-dimensional structure was not modeled for Rosβ, an AlphaFold model^44^ of the linear RosA1 precursor core peptide also predicts an alpha helix in the N-terminal region (Figure 5C), which is supported by the observation of (*i*, *i*+3) NOEs in the spectra of Rosβ. The directionality of ring formation is also revealed by the structure of Rosβ, indicating that all rings are formed from a C-terminal Cys that attacks a Dha/Dhb that is located N-terminal from that Cys (C-to-N direction of cyclization; Figure 5B). RosM is a remarkable class II synthetase in that it acts on two very different substrate sequences and makes two very different ring patterns. This finding contrasts with the two-component class II lanthipeptide synthetase CylM, which acts on two peptides with similar sequences and makes similar ring patterns (Figures 1C and S1). RosM also is different from the class II synthetase ProcM, which acts on 30 peptides with very different sequences,^45^ but catalyzes both C-to-N cyclization and N-to-C cyclization.^46–48^

**Figure 6.**
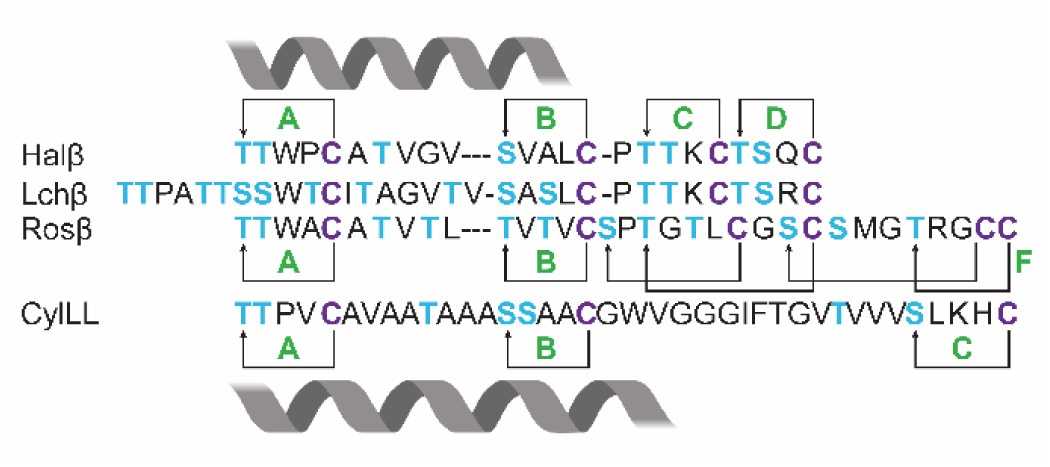
Structural similarity of Rosβ with other lanthipeptides. Created with BioRender (https://BioRender.com).

The unique structure determined here for Rosβ combined with Rosα lacking the Glu residue crucial for bioactivity described for other mersacidin-like peptides, supports a potential novel mode of action compared to other examined two-component lantibiotics that may not be lipid II dependent. The determination of the structure of Rosβ sets the stage for mechanistic studies on its mode of action.

## Experimental Section

### General materials and methods

Materials and reagents were purchased from Gold Biotechnology, Fisher Scientific, or Sigma-Aldrich unless otherwise noted. Molecular biology reagents for cloning (Q5 polymerase, and deoxynucleotides) were purchased from NEB. Oligonucleotide primers and genes were obtained from Integrated DNA Technologies and Twist Bioscience. DNA plasmids were prepared using MiniPrep kits (Qiagen) using the manufacturer’s instructions. Sanger sequencing was performed by the Roy J. Carver Biotechnology Center (University of Illinois at Urbana-Champaign), ACGT, and SNPsaurus. MALDI-TOF-MS analysis was performed using a Bruker UltrafleXtreme MALDI-TOF mass spectrometer (Bruker Daltonics) in reflectron positive mode at the University of Illinois School of Chemical Sciences Mass Spectrometry Laboratory. All samples were analyzed using Super-DHB MALDI matrix. Multiple sequence alignments were performed using ClustalW.

### Plasmid construction and generation

All cloning and mutagenesis experiments were performed using *E. coli* DH5α cells. The RosA1:RosM:pET28a vector to co-express RosA1 and RosM was reported previously.^22^ A previous gene construct for the expression of LahT_150_ in a pETDuet-1 vector with a His_6_-tag was also utilized for this study.^38^ Information for plasmids used or generated can be found in Table S1.

### Protein and peptide expression

All expressions were performed using *E. coli* BL21 Rosetta2 (DE3) unless otherwise specified. All constructs were overexpressed using TB medium. All cells transformed with the pET28a vector were grown in the presence of 50 µg mL^-1^ kanamycin, and those transformed with pETDuet-1 were grown in the presence of 100 µg mL^-1^ ampicillin.

Starter cultures were made for each expression using TB medium and were supplied with antibiotics indicated above and inoculated with a single colony. Starter cultures were incubated at 37 °C and shaken at 220 rpm overnight. Expression cultures were inoculated with 10 mL of starter culture per L of expression, supplied with antibiotic, and incubated at 37 °C with shaking (200 rpm) until an OD of 0.8-1.0 was reached. The culture was then cooled at 4 °C for 30 min, and IPTG was added to a final concentration of 1 mM to initiate over-expression. The cultures were incubated at 18 °C with shaking (200 rpm). Cells were harvested after 18 h and stored at −70 °C until purification.

### Protein and peptide purification

The modified RosA1 peptide was purified by resuspending cell paste in 20 mL of LanA B1 buffer (6.0 M guanidine HCl, 20 mM NaH_2_PO_4_, 0.5 mM imidazole, 1 mM TCEP, pH 7.5) for 1 L of cell culture and lysis via sonication at 60% amplitude for 5 min with a 2.0 s on and 6.0 s off pulse. The lysate was clarified via centrifugation at 50,000×g for 1 h and the supernatant was filtered using 0.45 µm syringe filters. The filtered lysate was loaded onto a gravity flow Ni-NTA column with 1 mL of resin preequilibrated with 6 column volumes (CV) of LanA B1 buffer (6.0 M guanidine HCl, 20 mM NaH_2_PO_4_, 0.5 mM imidazole, 1 mM TCEP, pH 7.5). The column was washed with 10 CV of LanA B1 buffer, 5 CV of LanA B2 buffer (6.0 M guanidine HCl, 20 mM NaH_2_PO_4_, 30 mM imidazole, 1 mM TCEP, pH 7.5), and 5 CV of wash buffer (20 mM NaH_2_PO_4_, 30 mM imidazole, 300 mM NaCl, 1 mM TCEP, pH 7.5). The peptide was then eluted using 10 CV of elution buffer (20 mM NaH_2_PO_4_, 1 M imidazole, 100 mM NaCl, 1 mM TCEP, pH 7.5). After purification of RosA1 that was modified by RosM *in vivo*, endoproteinase AspN was used to remove a large portion of the leader peptide. The buffer of the purified peptide was first exchanged into AspN digestion buffer (50 mM Tris HCl pH 8.0, and 2.5 mM ZnSO_4_; 2X reaction buffer provided by NEB) using a 3 kDa Amicon centrifugation filter and the sample was concentrated to approximately 5 mL. Digestion reactions contained a mass-to-mass ratio of 50:1 of peptide: AspN. Reactions were incubated for 2 h at 37 °C and AspN-digested peptides were purified via reversed phase HPLC using a Phenomenex Luna C5 semiprep column (250 × 10 mm, 10 mm, 100 Å) at a flow rate of 8 mL min^-1^ and with the following gradient over 32 min: 2% B for 10 min, 2– 30% B over 2 min, 30–60% B over 15 min, 60–100% B over 2 min, and hold at 100% B for 3 min (A: 0.1% TFA in H_2_O, B: 100% MeCN, 0.1% TFA). HPLC-purified peptide was collected and lyophilized, and the peptides were stored as a powder at −20 °C until use. Then additional residues of the predicted leader peptide were removed utilizing proteinase K as previously described to obtain Rosβ peptide.^8^

LahT150 was expressed and purified as previously reported. Its activity was confirmed with ProcA2.8 as substrate.^38^

### Structural characterization

For NMR studies, the Rosβ peptide was dissolved in 530-560 µL of 50% acetonitile-d_3_ in H_2_O and transferred to a 5 mm thin wall precision NMR sample tube (7” L, 535-PP, purchased from Wilmad-LabGlass). An ^1^H double pulsed field gradient spin-echo (DPFGSE) TOCSY with a 70 ms mixing time and DPFGSE NOESY with a 350 ms mixing time were acquired at 25 °C. The data were acquired on an Agilent VNMRS 750 MHz spectrometer using VNMRJ 4.2A software with the BioPack suite. The acetonitrile signal was referenced as 1.97 ppm prior to manual data analysis.

For the Rosα peptide, we tried the following solvent systems: acetonitile-d_3_ (MeCN-d_3_); MeCN-d_3_ in H_2_O, varying the fraction of MeCN (10-60%) and with or without 0.1% TFA; H_2_O/D_2_O with 0.1% TFA; MeOH-d_3_. In most cases, solubility was very poor. In solvent systems where solubility was somewhat better, signal dispersion was poor and we could not assign the amide signals, probably because of aggregation. We note that DMSO oxidizes the thioether crosslinks and cannot be used.

### Determination of stereochemistry

Stereochemistry determination was adapted from the previously reported advanced Marfey’s method.^39^ The peptide sample was hydrolyzed in 6 M DCl for 20 h at 120 °C. The resulting hydrolysate was derivatized with 1-fluoro-2,4-dinitrophenyl-5-L-leucinamide (L-FDLA) in 0.8 M aqueous NaHCO_3_ for 3 h at 67 ℃. Following the reaction, the mixture was neutralized with 6 M

HCl and lyophilized. The dried residue was then extracted with MeCN to remove residual inorganic salts. The extract containing the diastereomeric derivatives was analyzed by LC-MS using an Agilent 6545 LC/Q-TOF instrument with a Kinetex F5 Core−Shell HPLC column (1.7 μm, F5 100 Å, 100 mm× 2.1 mm). mCylL_s_ (plasmid available at Addgene ID #208760) and mCoiA1 (Addgene ID #208761 and #208762) were used for lanthionine and methyllanthionine standards, respectively, for comparison and co-injection.

## ASSOCIATED CONTENT

### Supporting Information

The Supporting Information is available free of charge at https://pubs.acs.org/doi/ Figures S1-S10 showing spectroscopic data, and Table S1 with gene sequences (PDF).

All NMR data has been deposited in: https://doi.org/10.57992/nmrxiv.p152

## Notes

The authors declare no competing financial interests.

