## Supporting figures and table for "An unusual ring pattern in the Rosβ lanthipeptide of the two-component lantibiotic roseocin"

Data associated with this study have been deposited at the link below and will be released upon publication:

Desormeaux, Emily; Zhu, L; van der Donk, Wilfred (2025), "An unusual ring pattern in the Ros-beta lanthipeptide of the two-component lantibiotic roseocin"", Mendeley Data, V1, doi: 10.17632/rnjfczgcw2.1

All NMR data has been deposited on NMRxiv: *https://doi.org/10.57992/nmrxiv.p152*

**Table of Contents**

Figure S1. Structures of two-component lanthipeptides that like roseocin lack the lipid II binding……S3
Figure S2. MALDI-TOF mass spectrum of RosA1 co-expressed with RosM and cleaved by endoproteinase AspN……S4
Figure S3. TOCSY spectrum of RosM-modified Rosβ ……S4
Figure S4. TOCSY-NOESY spectra overlay of RosM-modified Rosβ showing the amide region……S5
Figure S5. TOCSY-NOESY spectrum overlay of RosM-modified Rosβ used for A ring assignments……S6
Figure S6. TOCSY-NOESY spectrum overlay of RosM-modified Rosβ used for B ring assignments……S7
Figure S7. TOCSY-NOESY spectrum overlay of RosM-modified Rosβ used for C ring assignments……S8
Figure S8. TOCSY-NOESY spectrum overlay of RosM-modified Rosβ used for D ring assignments……S9
Figure S9. TOCSY-NOESY spectrum overlay of RosM-modified Rosβ used for E ring assignments……S10
Figure S10. TOCSY-NOESY spectrum overlay of RosM-modified Rosβ used for F ring assignments…… S11

Table S1. DNA sequences of genes used……S12

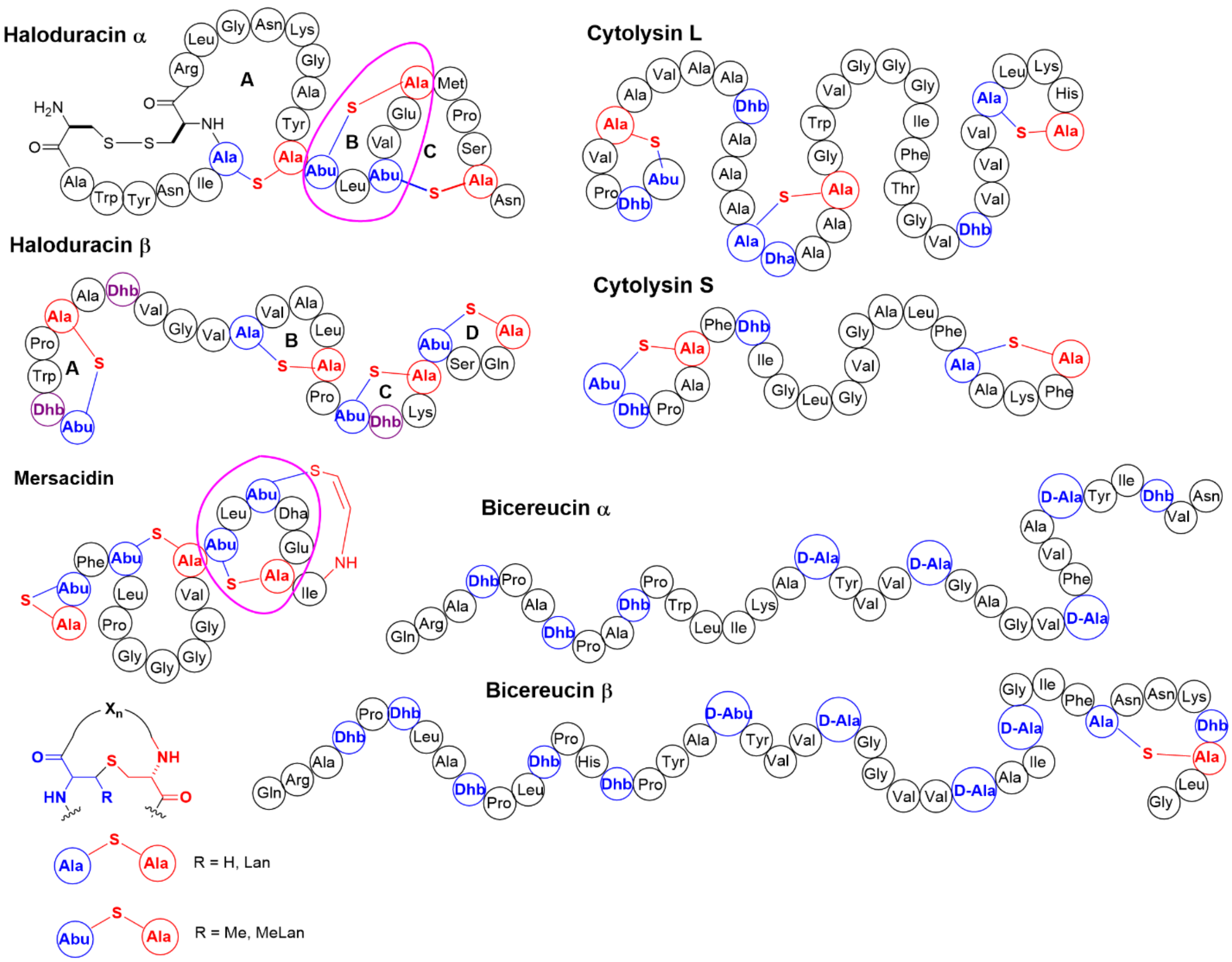


**Figure S1.** Structures of two-component lanthipeptides that like roseocin lack the lipid II binding motif of mersacidin. Haloduracin and mersacidin are shown for comparison with the lipid II binding motif circled in magenta. Structures derived from Ser/Thr are in blue, structures derived from Cys are in red. The shorthand notation used for Lan/MeLan is shown in the left bottom corner.

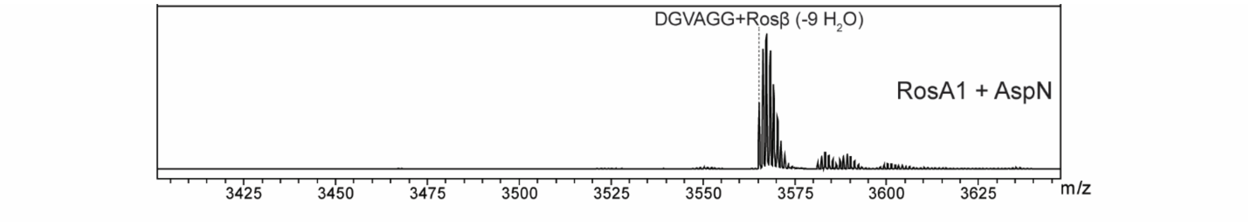


**Figure S2.** MALDI-TOF mass spectrum of RosA1 co-expressed with RosM and cleaved by endoproteinase AspN. This process leaves six residues at the N-terminus of Rosβ.

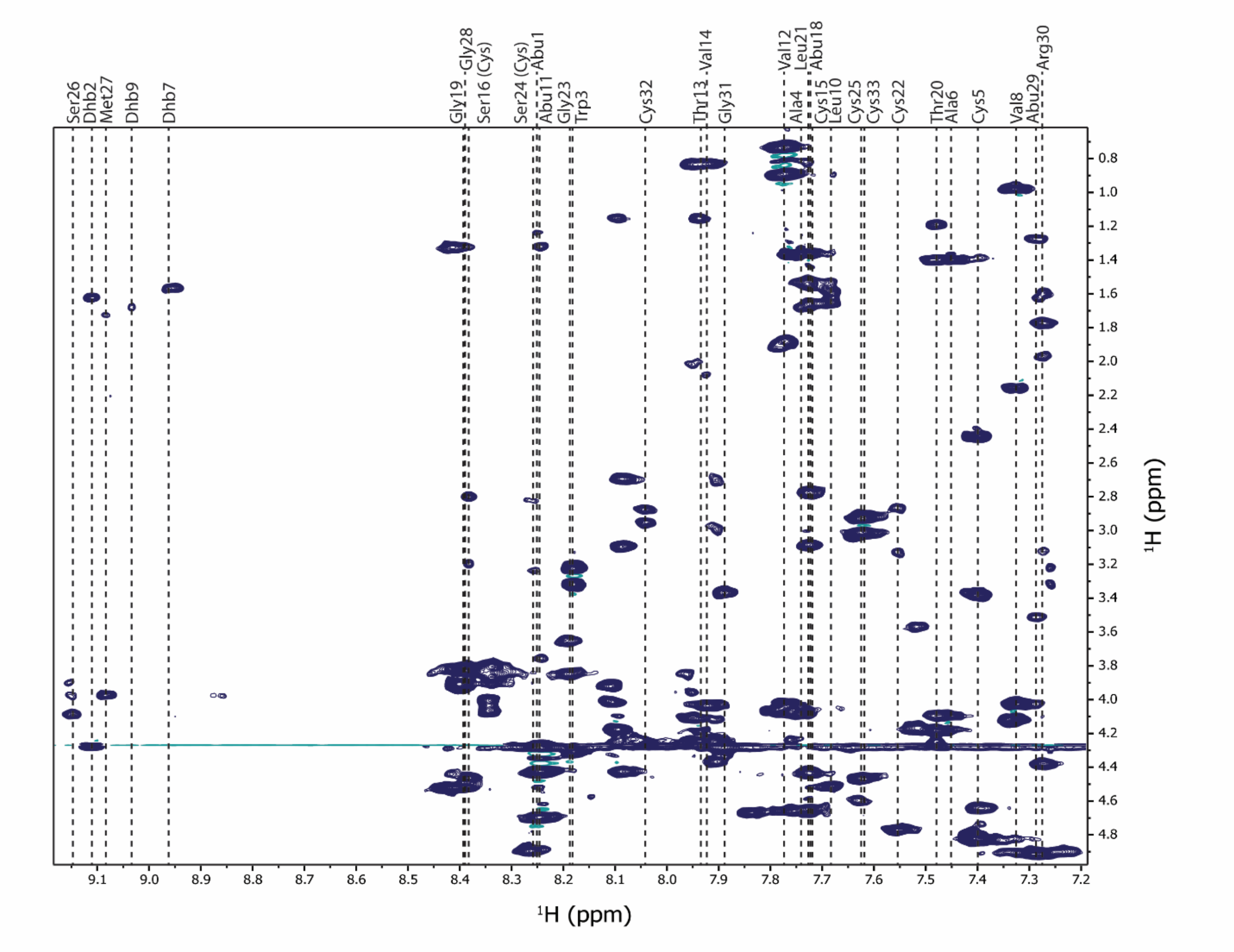


**Figure S3.** TOCSY spectrum of RosM-modified Rosβ in 50% ACN-$d_3$/50%$H_2O$ showing the spin system of its component residues without axis breaks.

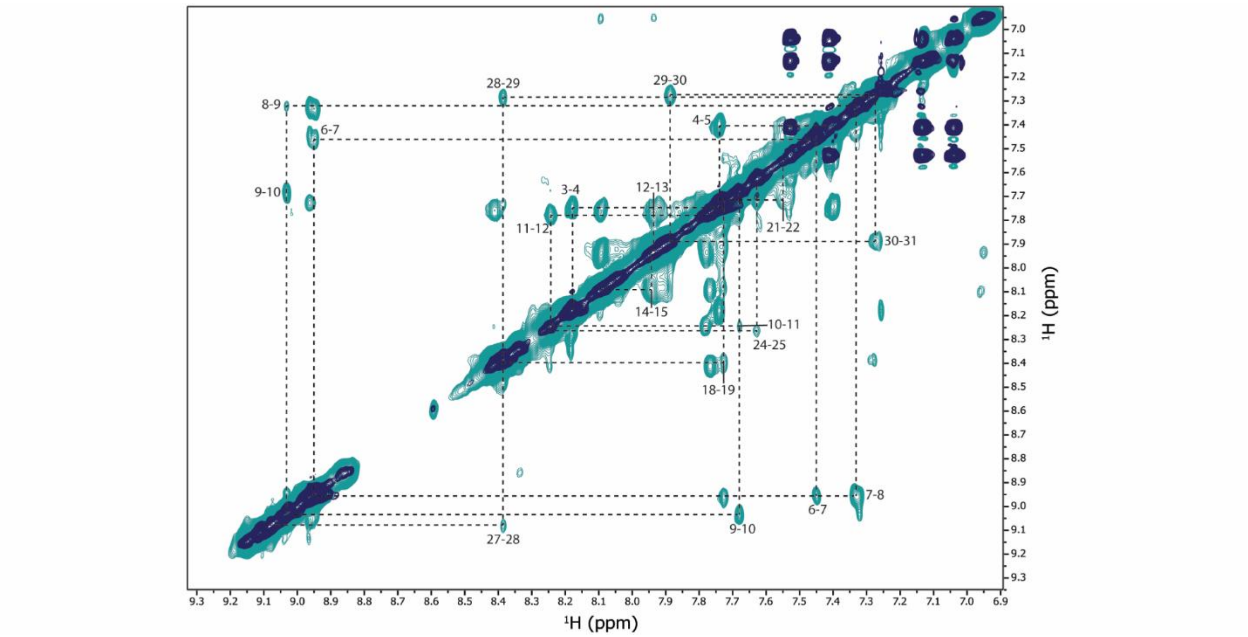


**Figure S4.** TOCSY-NOESY spectra overlay of RosM-modified Rosβ showing the amide region. Some amide resonance assignments are indicated. Some peaks with less intensity are omitted for figure legibility.

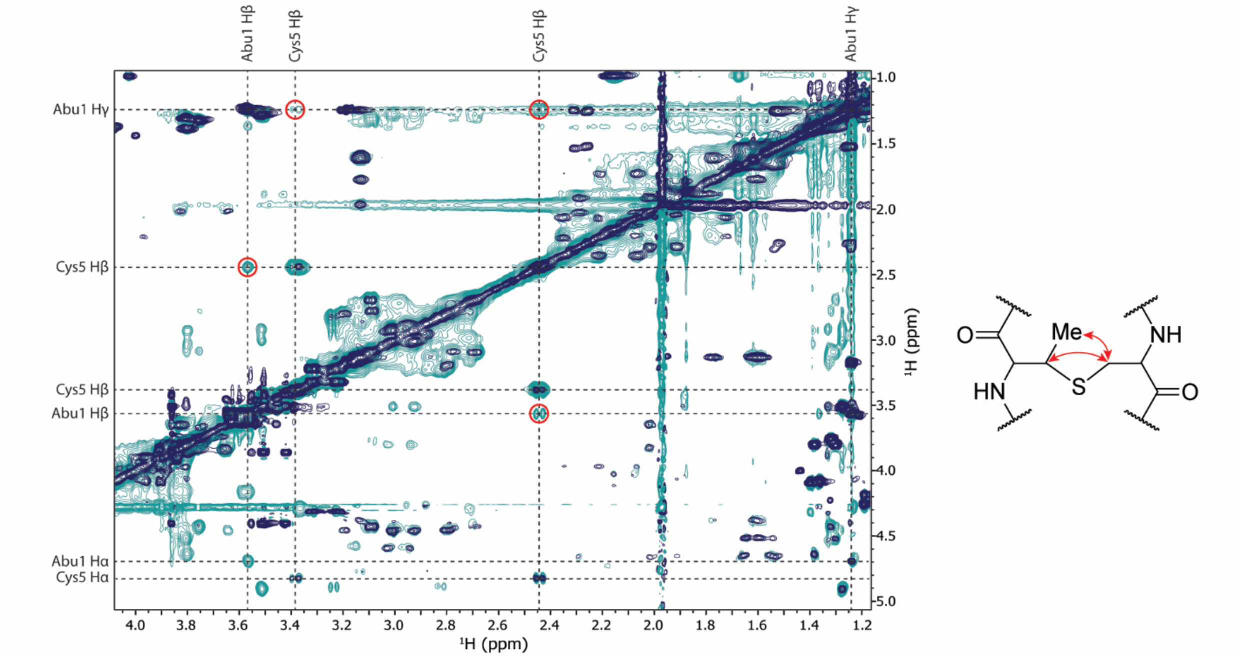


**Figure S5.** TOCSY-NOESY spectrum overlay of RosM-modified Rosβ used for A ring assignments. The TOCSY spectrum is indicated in dark blue, and the NOESY spectrum is indicated in cyan. NOESY cross peaks indicative of A ring formation between Thr1 and Cys5 are circled in red, and the corresponding couplings are indicated on the MeLan ring drawn to the right with red arrows.

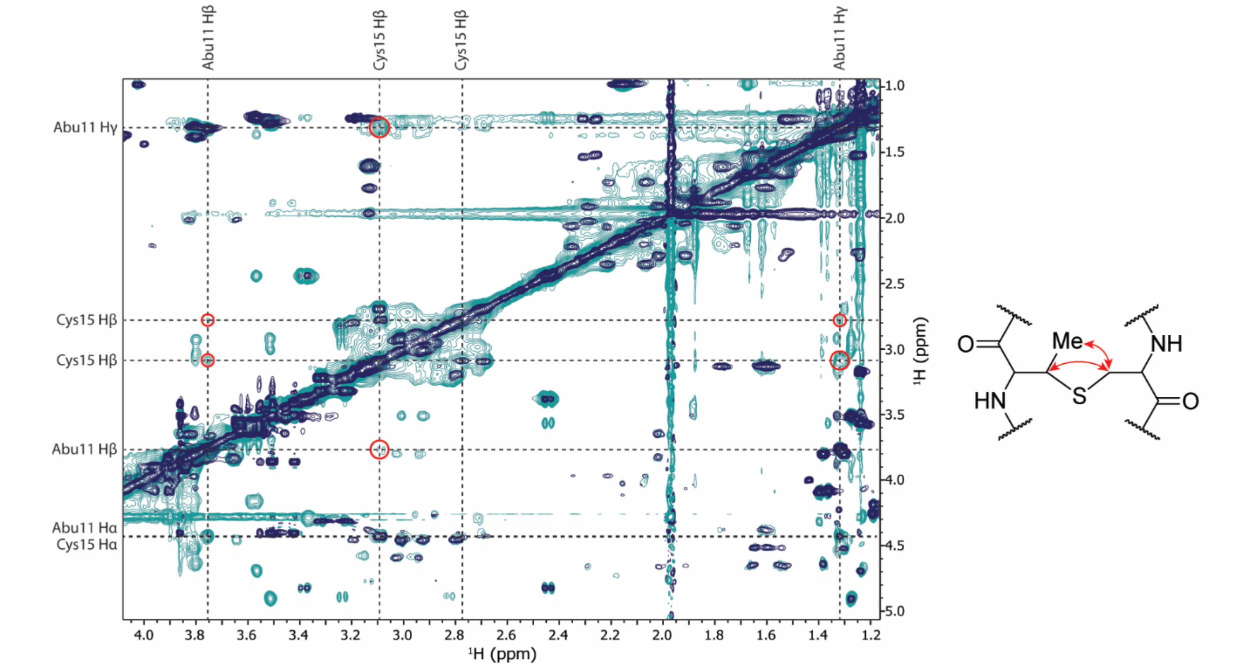


**Figure S6.** TOCSY-NOESY spectrum overlay of RosM-modified Rosβ used for B ring assignments. The TOCSY spectrum is indicated in dark blue, and the NOESY spectrum is indicated in cyan. NOESY cross peaks indicative of B ring formation between Thr11 and Cys15 are circled in red, and the corresponding couplings are indicated on the MeLan ring drawn to the right with red arrows.

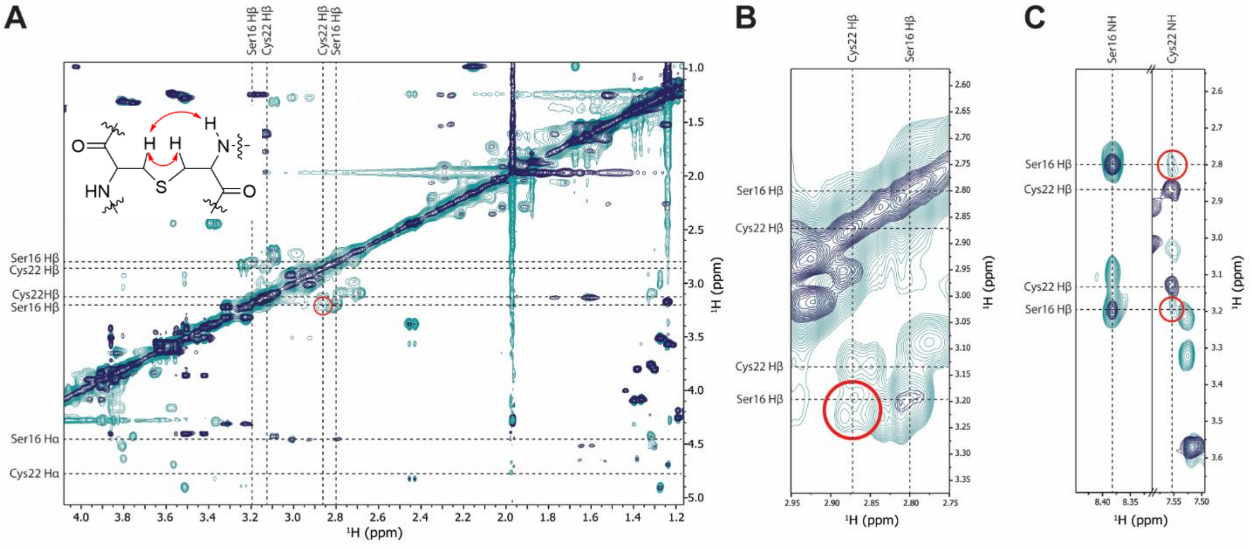


**Figure S7.** TOCSY-NOESY spectrum overlay of RosM-modified Rosβ used for C ring assignments. The TOCSY spectrum is indicated in dark blue, and the NOESY spectrum is indicated in cyan. NOESY cross peaks indicative of C ring formation between Ser16 and Cys22 are circled in red, and the corresponding couplings from panels A-C are indicated on the Lan ring drawn with red arrows. **A)** Spectrum overlay showing the aliphatic region. **B)** Zoomed in view of the NOE peak between the β protons of the Lan forming residues. **C)** Spectrum overlay showing the cross peaks between the Cys22 amide proton and the Ser16 β protons.

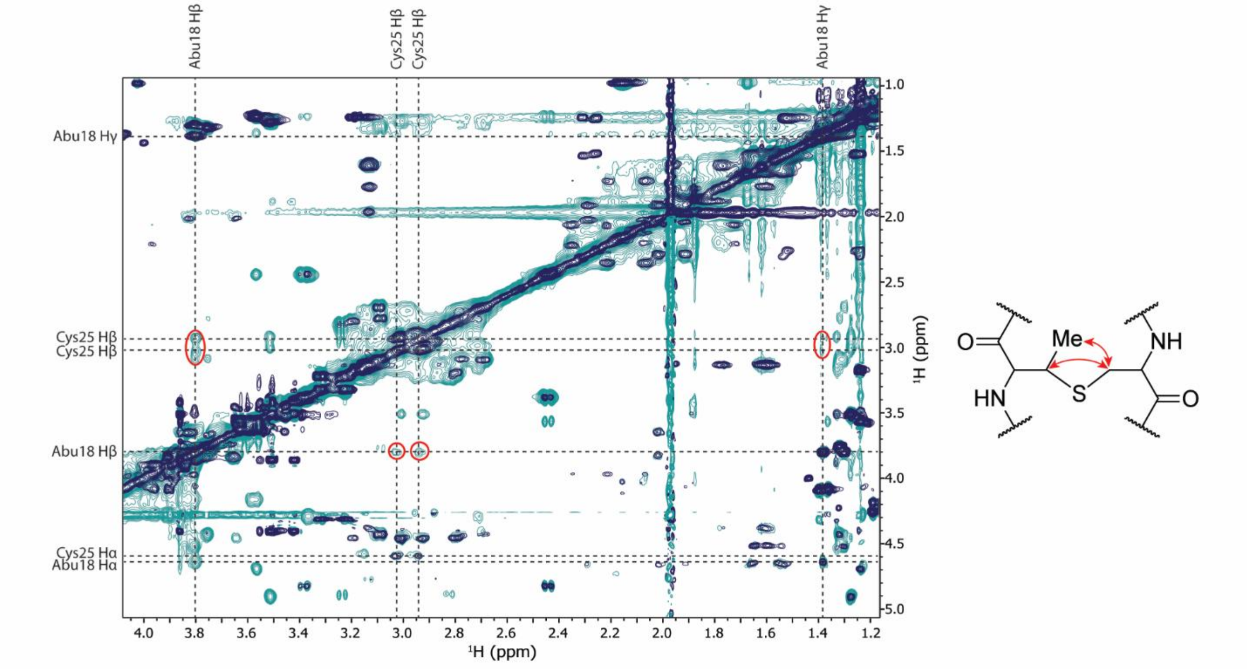


**Figure S8.** TOCSY-NOESY spectrum overlay of RosM-modified Rosβ used for D ring assignments. The TOCSY spectrum is indicated in dark blue, and the NOESY spectrum is indicated in cyan. NOESY cross peaks indicative of D ring formation between Thr18 and Cys25 are circled in red, and the corresponding couplings are indicated on the MeLan ring drawn to the right with red arrows.

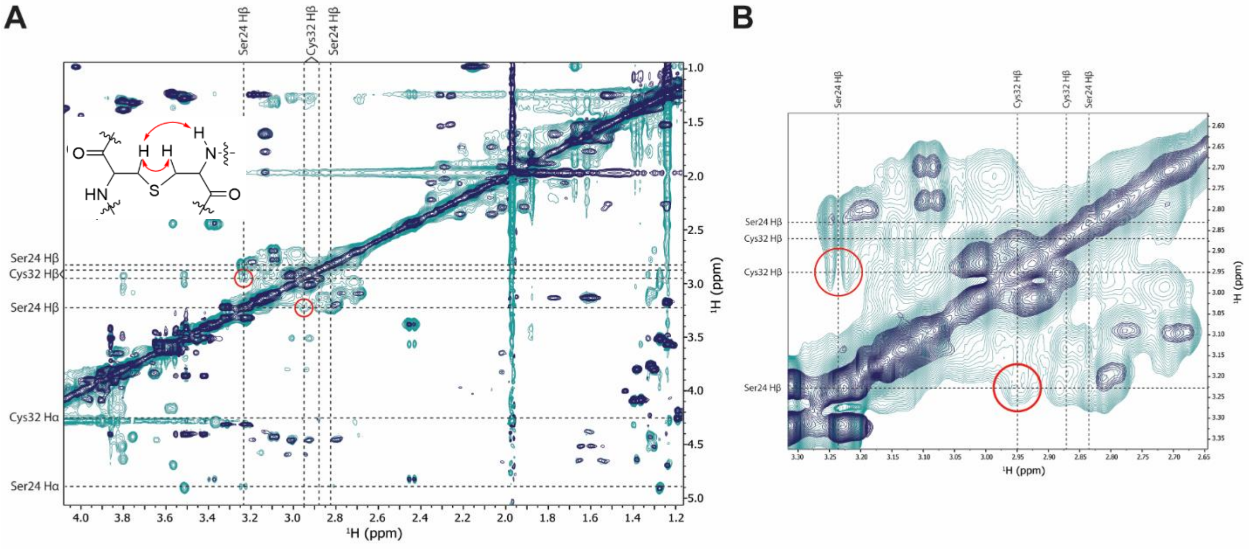


**Figure S9.** TOCSY-NOESY spectrum overlay of RosM-modified Rosβ used for E ring assignments. The TOCSY spectrum is indicated in dark blue, and the NOESY spectrum is indicated in cyan. NOESY cross peaks indicative of E ring formation between Ser24 and Cys32 are circled in red, and the corresponding couplings from panels A and B are indicated on the Lan ring drawn with red arrows. **A)** Spectrum overlay showing the aliphatic region. **B)** Zoomed in view of the NOE peak between the β protons of the Lan forming residues.

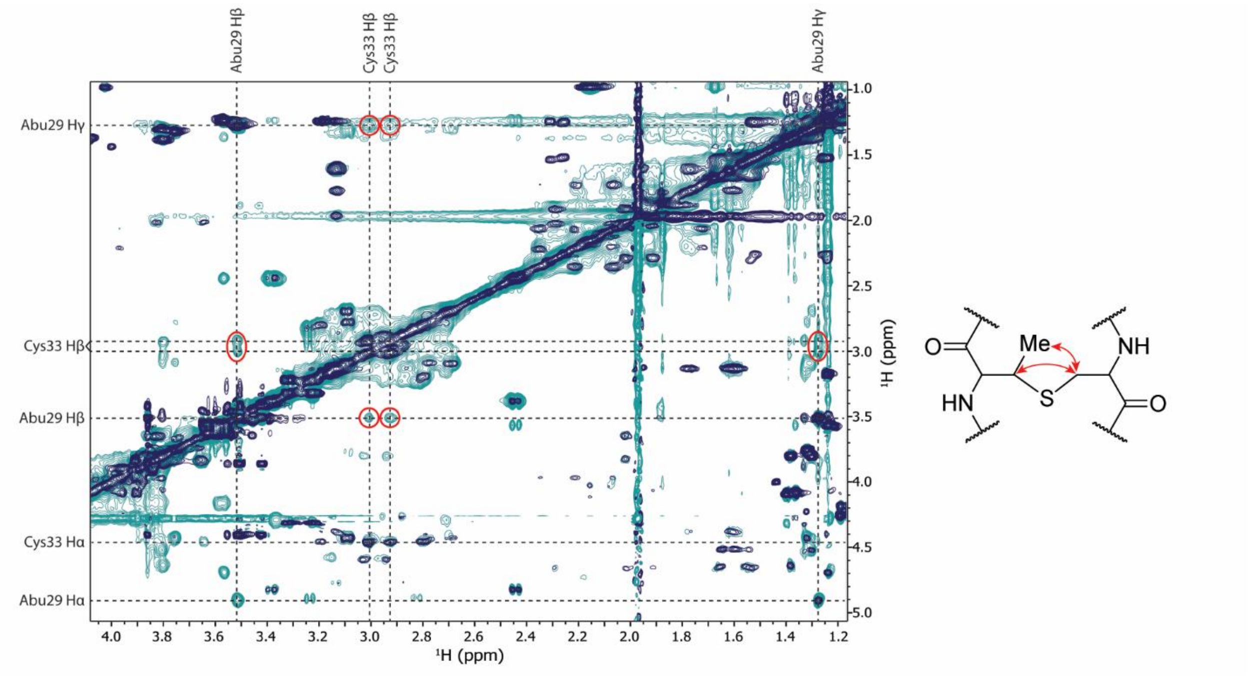


**Figure S10.** TOCSY-NOESY spectrum overlay of RosM-modified Rosβ used for F ring assignments. The TOCSY spectrum is indicated in dark blue, and the NOESY spectrum is indicated in cyan. NOESY cross peaks indicative of F ring formation between Thr29 and Cys33 are circled in red, and the corresponding couplings are indicated on the MeLan ring drawn to the right with red arrows.

**Table S1. DNA sequences of genes used.** All sequences were codon optimized for expression in *E. coli*.

| Protein/Peptide Name | Sequence |
|---|---|
| $His_6$-$LahT_{150}$ | ATGGGCAGCAGCCATCACCATCATCACCACAGCCAGGATCCGAGTAAAAAGCAGATACAGCCTGTCACAAGAGGGCG<br>AGCCAAGGTTCCTGTTATCATGCAGATGGAAGCTCTCGAATGTGGCGCGGCATCACTTGCAATGGTGCTTGCTTATT<br>ATAAAAAATGGGTACCGCTCGAGCAGGTCAGAGTTGACTGCGGTGTATCTCGTGACGGTTCCAATGCACTGAATGTG<br>CTTAAGGCAGCACGAAACTACGGTCTTGAAGCGAAGGGTTATCGTTATGAACCGGAGAAGCTGAAGAAGGAAGGCAC<br>CTTCCCGTGTATTATACACTGGAATTTTAATCATTTTGTTGTTCTTAAAGGTTTCAAGGGGAAATATGCATATATAA<br>ACGATCCTGCAAAGGGTGATGTTAAAATACCGATGGAGGAGTTTGATCGTTCTTTTACGGGTATATGTCTCATTTTC<br>AAGCCTACTGATAGATTTGAACAGTAA |
| RosM | ATGCCAGATGATGCTTCCGAACATCCTACTGTACGTCCTGATGGTGTGCATCTGCCATGGCACCGTGCAGCAACGCT<br>GTCAGAACGTTTGGGTGGAACACCATCGCGTTCGGATCCAGTTCGTGGTGAGAAACGCTTGGCGATGTGGCGTGAGC<br>TTGCTCCGCTCACGGGTGAACGTGCCACCTTAGATGCACGTTTAGCTCCGCTTGGTGTTGATGCGGCGGAACTGACC<br>GCGTTACTTGGCGAGTCAGATGATAGTCTTGCAGCCCGTACCGCCGACGAACCGGATTGGCATCGTACCTTTCGTCG<br>CTGGTGGGCAGAAGGGATGGCCACTACGGATGCAGCCCGTCGTTTGGATCCGGATATGGGCTTGCTTGAAGTGATTC<br>GTCCGCTGGTCGAAGGAGCCGTACATGAATTGGATGCTCTGGTGGAAGAACGCGTTCGTGCGTCCGATGATCCTGTA<br>ATGGCTGATCCCGGGCATATTCTGGACGTCGTCCGTGCGACTGCCCCCGCAGCGGATCTCTTGGGTCATGTTCAACG<br>TACGTTAGTTCTTGAACTTAACATTGCCCGTGTTGAGGAACGTTTAGGTGGTGAGACTCCTGAAGAACGTTTTAGTT<br>CCTATGTCGAGTTGCTGCGCAGCCAGGGTAACGCATTAGCATTGTGGGATGAATATCCCGTGTTAGCTCGTTTGGTG<br>GTTAATCAGCTTCGCTTTTGGATTGATACCCGTGTTGAACTGATCGATGCCCTGGTAGCCGATTTGCCAGCCTTGCG<br>CGGTGGGCTTCTCCCGAGCCCAGGGCCACGTCGTCTGGAACGTTTAGAATTTGGGGCGGGTGATAAACATCGTCGTG<br>GCCGTAGCGTGGCTATGGTAGAATTTGATACCGGAACCCTGGTGTTTAAACCACGTAGTTTAGCGATGGATACGGCG<br>TTTGATGGTTTGCTGGCATGGGTAAACCGTCAACAACCCCCACATGATCTGGCACGCATTGGTGTGTTAGAACGCGG<br>CGGCCACGGCTGGGTTGAACAAGTAGATACCACGGCCACCACCGATGCTGCCGGTGGTGATCGTTATGCATGGCGTC<br>TCGGCGCACTGACCTCACTGTTATACTTACTTCACGCGACCGATTTTCATTTTGAAAATGTTTTGGCGGCAGGTGAA<br>CATCCCGTCTTGGTTGATCTGGAAGCACTTTTACATAACGATAAAACTGCCGCAGTTGTTAAGATTGATGGAGAGAC<br>TGATATTGCAGCCGTTACCCTGGCGGATAGTGTACAAAGTGTAGGAATCTTACCGAATCACTTGTTAGTACGTGGCG<br>AAGATGGCACGTTTGGTCTGGATGTTTCAGGATTGGCAGGCCACGGTGGCCAATTAACGCCTATGCCCGTGCCGACC<br>TGGGAAGATTCTGGCACTGATCGTATGCGTCTCGTGAAACGCCGTATGGAAATGGATGCAGAACGTAACCACCCGAT<br>GACCGCCGATGGCGAACCCATTGACCTCGTTGGTCGTGTGGACGCATTTGTCGATGGGTTTACACACGTTTACCGTC<br>TCCTGGAAGCCGGCCGTGATGAGCTTCTGGATCCGGCTGGTCCTTTGGCAGCCTTTGGCTCGGTGCGCACCCGTTTA<br>ATTGCTCGCCCAACCCATATTTATGGTCGTTTATTACTCGAGTCAACGCATCCCGATTTCTTGCGTGATGGGTTAGA<br>TCATGCACGTAGTCTTGCGCGCCTGGCGGGCGGACATGTAGAAATGCCGGAATCTCGTGAAGCCATGGTTCAGGATG<br>AGGTAGCAGAATTGACACTCGGAGATATCCCGATGTTTACCGTGGACGCAGAAAGTGGCACCTTGCGTGGCGGTTTT |

GATGGCCGTCCAATTGGAACTCGCTATCCACCGTTAGCGGCAGTACGCGAACGTTTAGCCGCCTTGGGCGATGCTGA
TCTGGCATTTCAGGAACGCGTTATTCGTTCATGTGTCGCAGCGACCGCCATGGGTGATTCGGACGCGCGTTGGCCTA
ATTGGCATCGTCCTCGCCTGGATGGTGCAGCAGACCCGGGTGACTTTGCAGAAGAAGCGCTGCGCTTAGCACGCCGT
TTAGGCGACTTATGTGTACGTGATGAACATGGTCTTGGCTGGATTGGATTGGATCTGGTTGATGAAAAGTATTGGCA
ATACGCGCCGGCCCCTATTGGGCTTTATACCGGTACGGCAGGAATTGCGCACGCGTTAGATGCGGTTGCAGCCGTAA
CGGGAGATGAAGCTACCGGGGAACTGGCACGCGCGGCATTCGATCAAGTAGCCAAACGCTCAGTATTAATTGCGGAA
ACCCTCGCTAGCTTAGCAAAGAAACCTGGCGCTGCGGAATTCGGTATTGGTGCGTTTGGCCCGTTTGGTGGTGCAGT
GTATGCCTTGGCTCATGCCGCGGTGCGTCATGATCGTCCTGATTATGCAGAGGCTGCAGCAGCATTACTCCCTGCAA
TTGATGAATTAGTAGAAGAGGACCCCCTGCTTGACGTTGTGTCGGGTTCAGCCGGTGCTATTCTGGCTCTGCTGGCT
CTTGAGAAAGCACGTCCTGCAGCGGGTGCTGCTCGCATTGCGGCTCGTTGTGCGGAGCGTCTGCTGGCTACACGTCA
AGAATGTGCAGAGGGCTGGGGTTGGGCAACGCCTATTAACCCTGAGGCTCCCTTGGCAGGCTTCAGCCATGGTGCGT
CAGGGATCGCATATGCCCTTGCGCGTTTAGATGCGGTGGCGCCGCGCCCAGAATATGGCGAAGCGGTGCGTAACGCC
CTGCGTTACGAACGTACGGTATTTGATCCTGAATTACGTACTTGGCGTGATCTTCGTCCGGCAAATCTGGGTGGACG
CACTGTAATGAACGCCTGGTGTCACGGGGCGCCCGGCATTGCGCTGGCCCGCGATGCTTTTCAGGGTTTAGGTACGG
CGGCCGATCTGGCAGATTTGGTTGAAAGCGATCGTCATACTGCCGTTTTGGCAGCTGTTAGCACGGGTTTAGATCTG
GATCCAGTGAGCGGGCTGGGTAATCATAGCTTATGTCACGGGGATGTTGGTAATTTATTAATTATCGAAGCAAGCGC
GCGCGCAGATCGCGAACCGGAAGTTGCGGGTTTGTTGCCACGTGTGTGGAATACCCTGCTGCATGAGGGCCGTGAAA
ACGGCTGGCTCTGTGGTGTCCCTAAAGGAGTGGAAACTCCAGGGTTAATGACCGGCTTAGCTGGCATCGCTTGGGGA
CTGGCACGTAAGGCGGCACCTGAACGTGTTCCGGATCTGTTGACCTTGGCGGCTCCAGCAGGACCTGGCGGGACCCC
CTAA
